# Reprogramming method does not impact the neuronal differentiation potential of 16p11.2 deletion patient iPSCs

**DOI:** 10.1101/2023.07.07.548182

**Authors:** Michael F. Wells, Ellen J. Guss, Hongyan Zhou, Bruce Sun, Hector Martinez, The NYSCF Global Stem Cell Array Team, Veronika Akopian, Scott Noggle, Daniel Paull, Jennifer Moore, Michael Sheldon, Julia E. Sommer, Marta Benedetti, Alexander Meissner, Kevin Eggan

## Abstract

A major impediment to the actualization of the induced pluripotent stem cell (iPSC)-based personalized medicine revolution is the lack of widely accepted standard operating procedures (SOPs) across different groups and institutions. The various methods employed can include choice of starting materials, reprogramming agents, and culture conditions, with each of these factors hypothesized to influence the reprogramming efficiency and transcriptional identity of iPSCs. As such, we systematically compared iPSC reprogramming procedures using cells derived from the somatic cells of three patients with 16p11.2 deletion syndrome (16p11.2del) and found remarkable similarity among the different methods. FACS analysis revealed that regardless of somatic cell type (fibroblast, lymphocyte, erythroblast), route of reprogramming factor introduction (mRNA, Sendai virus, episome), donor sex, or facility (Rutgers, NYSCF), 16p11.2del patient iPSCs were viable as high purity cultures expressing pluripotency marker proteins. This observation was supported at the transcript level by qPCR analysis, which demonstrated the ability for the iPSCs to differentiate into all three embryonic germ cell lineages after 12 days in culture as embryoid bodies. NGN2-mediated differentiation of these iPSCs produced functional neurons that formed active synaptic networks as revealed by multi-electrode array (MEA) recordings. Importantly, no group-wise comparisons among the reprogramming methods yielded consistent statistically significant differences, indicating that these procedures are equally capable of producing pluripotent stem cells that can efficiently differentiate into mature, functional neurons. This work highlights the utility of these reprogramming methods and supports the use of differentially reprogrammed iPSCs for direct comparative studies of human neurodevelopment.

## INTRODUCTION

Human induced pluripotent stem cells (iPSCs) have emerged as a groundbreaking technology with immense potential to revolutionize biomedical research and regenerative medicine. This approach involves the reprogramming of somatic cells, such as skin or blood cells, into pluripotent stem cells through the introduction of specific transcription factors. The resulting iPSCs hold the capacity to differentiate into any cell type in the human body, thus providing a powerful tool for modeling human biology in a dish. The utility of iPSCs extends beyond basic research, encompassing applications in drug discovery, disease modeling, and personalized therapies. Though this technology is primed to serve as the foundation of future personalized treatments, it is not without its technical shortcomings.

The iPSC reprogramming process has been shown to introduce novel mutations and epigenetic states not found in somatic donor cells^1–3^. In addition, early studies utilizing iPSCs have been plagued by high line-to-line variability, most likely due to inconsistencies in reprogramming efficiency^4–6^. Widely accepted standard operating procedures across different groups and institutions do not exist at this time^7^. Currently, numerous methods are used to reprogram somatic cells to iPSCs, with differences in starting material (e.g. fibroblasts, lymphocytes, erythrocytes), reprogramming agents (e.g. synthetic mRNA, episomal DNA, DNA viral vectors), and culture conditions. Given that each of these factors are suspected to influence the efficiency of reprogramming and the transcriptional identity of iPSCs, it is critical to systematically compare the efficacy and reproducibility of the methods currently used throughout the stem cell field.

To this end, we conducted a series of experiments aimed at directly comparing iPSCs reprogrammed via different procedures used at multiple institutes and service providers. We graded methods and providers using standard biochemical and functional assays with a particular focus on their ability to generate iPSCs that meet the basic requirements for pluripotency and display acceptable induction efficiencies into mature functional neurons.

All iPSC lines were derived from patients harboring 16p11.2 microdeletion (16p11.2del), which accounts for 0.5% of autism spectrum disorder (ASD) cases^8^. In addition to ASD, patients suffering from 16p11.2 deletion display a range of symptoms and diagnoses, including abnormal head size, intellectual disability, and developmental delay^8–10^. Together, the clinical manifestation and the prevalence of this large copy number variant in individuals with ASD suggest that this region contains genes that are critical for normal brain development. Though animal models of 16p11.2 rearrangements have provided valuable insight into the mechanisms underlying disease^11^, progress towards the discovery of effective clinical treatments has been hampered by the lack of detailed mechanistic studies using patient brain tissue. Induced pluripotent stem cells provide a potential means to circumvent this obstacle by affording researchers the opportunity to study neurodevelopmental disease using patient-derived neurons.

Remarkably, we found little to no differences across the 16p11.2del patient iPSC lines using our metrics, suggesting that each of the tested reprogramming methods are suitable for functional investigations of human neural cells. We did, however, discover abnormal network firing dynamics in the 16p11.2del patient lines relative to neurotypical controls using multi-electrode array electrophysiological recordings, thus highlighting a potential disease-related phenotype. As a whole, our results could not only guide decisions related to the establishment of high-throughput iPSC production standard operating procedures, but also influence the design of ASD-centric experiments that involve human cell types.

## RESULTS

### Description of 16p11.2del patient lines for comparative analysis

To effectively compare different reprogramming methods, we needed to test iPSCs derived from multiple patients by multiple facilities using multiple techniques. To this end, we received a total of 47 iPSC lines derived from three 16p11.2del patients (14758.x3, 14799.x1, 14824.x13) by two independent groups (NYSCF: New York Stem Cell Foundation Research Institute and RUCDR: Rutgers University Cell and DNA Repository). Different reprogramming starting materials (fibroblast, lymphocyte, and erythroblast) and reprogramming agent delivery systems (mRNA, Sendai virus, episomal DNA) were used to generate these iPSCs so that a total of 5 groups emerged for functional assessments: NYSCF-Fibroblast-mRNA (NFM), NYSCF-Fibroblast-Sendai (NFS), RUCDR-Erythroblast-Sendai (RES), RUCDR-Fibroblast-Episomal (RFE), and RUCDR-Lymphocyte-Sendai (RLS). Each group was represented by either two (RLS, RES) or three donors (NFM, NFS, RFE), and multiple clones were available for a majority of the patient lines within each group (Figure 1a).

**Figure 1:**
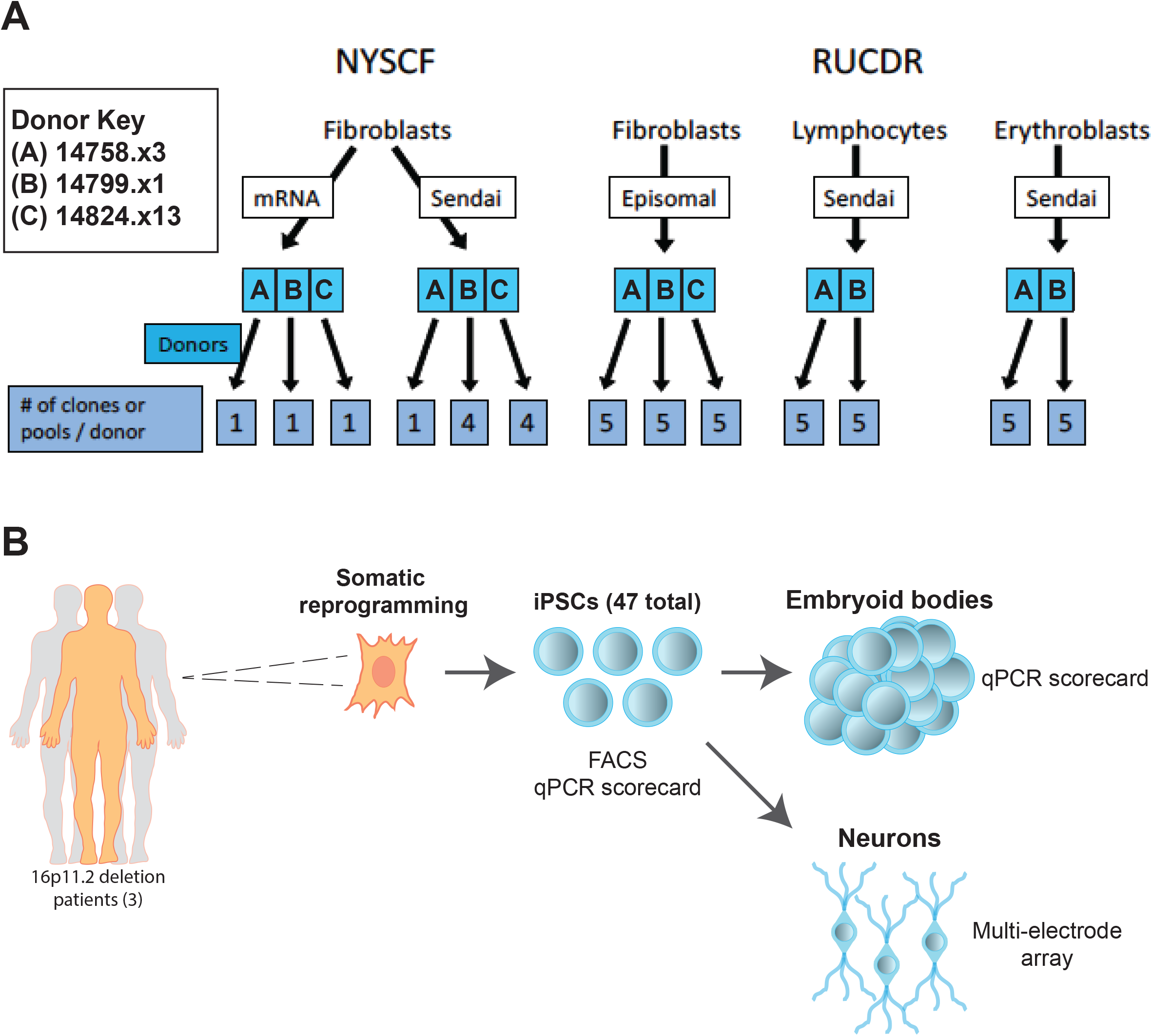
Project overview. **A**, Breakdown of 47 iPSC lines and the reprogramming methods used. **B**, Schematic describing overview of this comparative study.

There are two important considerations when assessing the results of our comparative analysis. First, we tested different numbers of clones per cell line due to the limited availability of some clones. This shortcoming inhibited properly powered comparative analysis for some of our assays. Second, NYSCF lines were polyclonal whereas RUCDR lines were monoclonal. Our design does not have the ability to discern the impact of these clonal differences on iPSC performance.

Cell line identities were coded by the service providers prior to delivery of live cells so that experimenters were blind to donor, method, and group throughout the duration of functional assessment. Upon receipt, cell lines were imaged for adequate confluency and colony morphology, expanded, and tested for mycoplasma contamination. Any cell lines not passing these initial quality control (QC) metrics were re-provided. Importantly, all 47 lines passed QC after the initial (37/47) or second delivery (10/47) of live cells from the respective group. Having received, expanded, and banked the 16p11.2del iPSC lines, we began our initial characterization of the 47 cell lines (Figure 1b). While we included iPSC and/or human embryonic stem cell (ESC) controls to benchmark our results, we ultimately based our conclusions on whether or not we saw consistent and biologically relevant differences among the reprogrammed 16p11.2del cells.

### Reprogrammed iPSCs express transcript and protein markers of pluripotency

Stem cells capable of differentiating into the three germ layers (ectoderm, endoderm, and mesoderm) tend to express high levels of pluripotency protein markers, such as SOX2, OCT4, and NANOG. To determine if reprogramming methods significantly affect expression of pluripotency markers, we compared the percentage of cells expressing these proteins across the 47 cell lines using intracellular immunostaining coupled with flow cytometry. Human ESCs and neurotypical (NT) iPSC controls were used as positive controls. As expected, a high percentage of the hESC and control iPSC cells expressed SOX2, OCT4, and NANOG (Figure 2). The 16p11.del iPSCs also showed consistently high levels of the pluripotency markers (Figure 2a). While statistical differences were sporadically identified between reprogramming groups, no difference was consistently discovered across all three markers (Figure 2a). The same could be said for analysis conducted on the data grouped by method of reprogramming agent delivery (Figure 2b), starting cell type (Figure 2c), donor sex (Supplemental Figure 1a), and patient (Supplemental Figure 1b). We did observe a minor difference when data was organized by genotype and facility (Supplemental Figure 1c-d). The NYSCF lines showed a slight (2-5%) decrease in the percentage of marker-positive cells compared to hES controls. Importantly, the percentage of marker-positive cells was at least 90% for both NYSCF and RUCDR lines, and no consistent differences were found between these two groups. It should be noted that the statistical difference between NYSCF and controls could be the result of the genotypic abnormalities inherent to the 16p11.2del lines or possibly the product of the dramatic differences in sample size between the two groups (16p11.2del: n = 47; NT: n = 11). These results suggest that, while the 16p11.2del lines may show higher variability in pluripotency marker expression compared to a small number of NT controls, the reprogramming method has no significant impact on the pluripotency protein expression of the resulting iPSCs.

**Figure 2:**
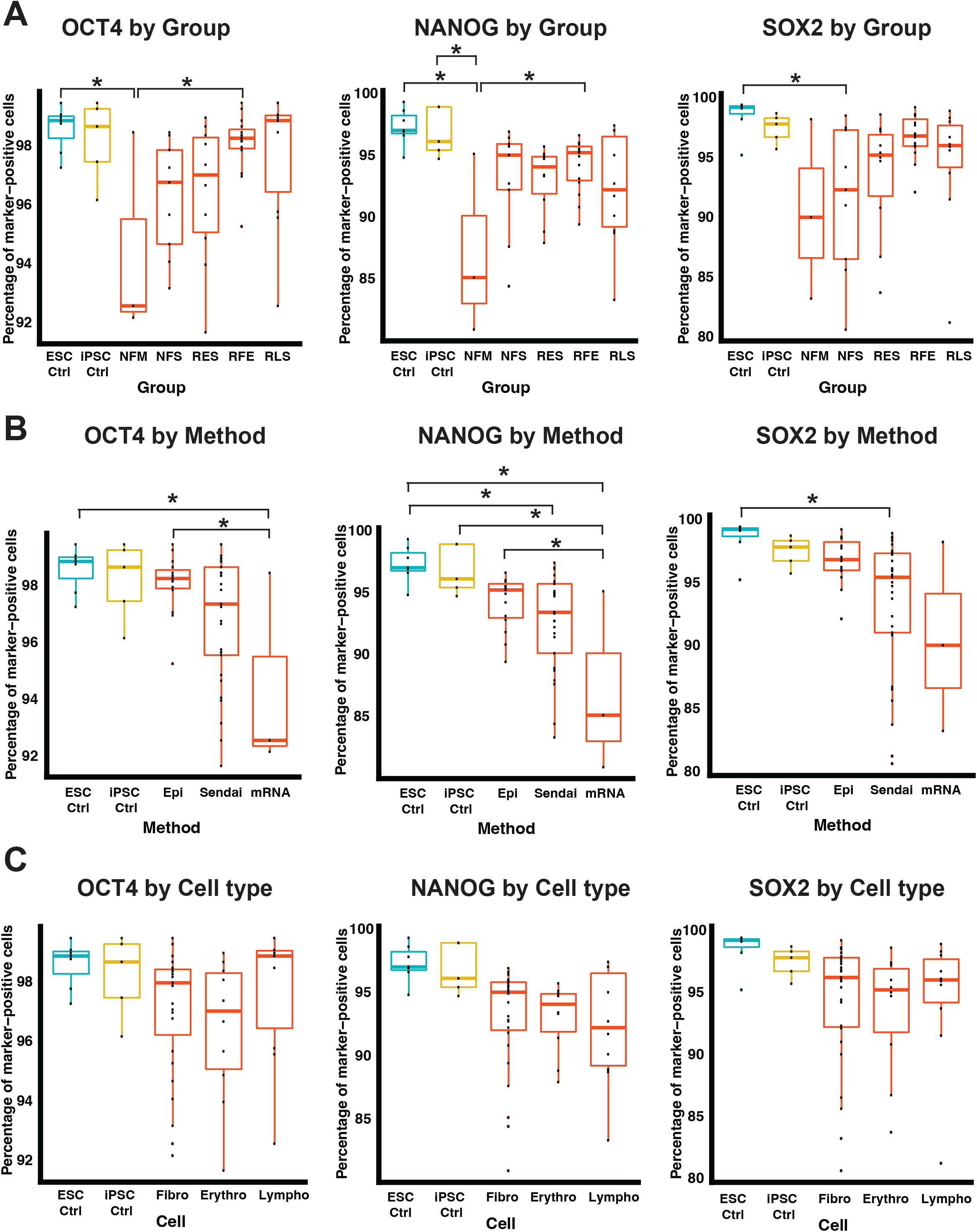
iPSC FACS analysis. **A-C**, Percentage of OCT4-, NANOG-, and SOX2-positive cells as a function of (A) reprogramming group, (B) factor delivery method, and (C) starting cell type. One-way ANOVA. *p<0.05.

While the protein expression of a handful of pluripotency genes is informative, we sought a more expansive and high-throughput way to quantify iPSC performance. We therefore employed the commercially available TaqMan hPSC Scorecard Panel, which is composed of 94 qPCR assays that measure a combination of control, housekeeping, self-renewal, and lineage-specific genes (ectoderm, endoderm, and mesoderm). A self-renewal score for each cell line was then calculated from the expression of nine pluripotency transcripts, where a range of +1 to -1 is considered normal. No differences were observed among the five 16p11.2del reprogramming groups, though the mean self-renewal score of the RLS group was significantly decreased compared to NT controls (Figure 3a). Small but significant differences were observed when 16p11.2del lines were compared to controls and grouped by delivery of reprogramming factors (Sendai vs controls; Figure 3b), starting cell type (Lymphocytes and fibroblasts vs controls; Figure 3c), reprogramming facility (NYSCF and RUCDR vs controls; Figure 3d), donor sex (Supplemental Figure 2a), and patient (14758.x3 vs controls; Supplemental Figure 2b). It is likely that these differences are primarily driven by differences in genotype, as shown by the NT control versus 16p11.2del comparison in Supplemental Figure 2c. For all sets of analysis, the mean self-renewal lineage score for each group fell within the normal range. Combined with the FACS analysis, these findings support the argument that the reprogramming method does not impact the pluripotential of the 16p11.2del iPSCs.

**Figure 3:**
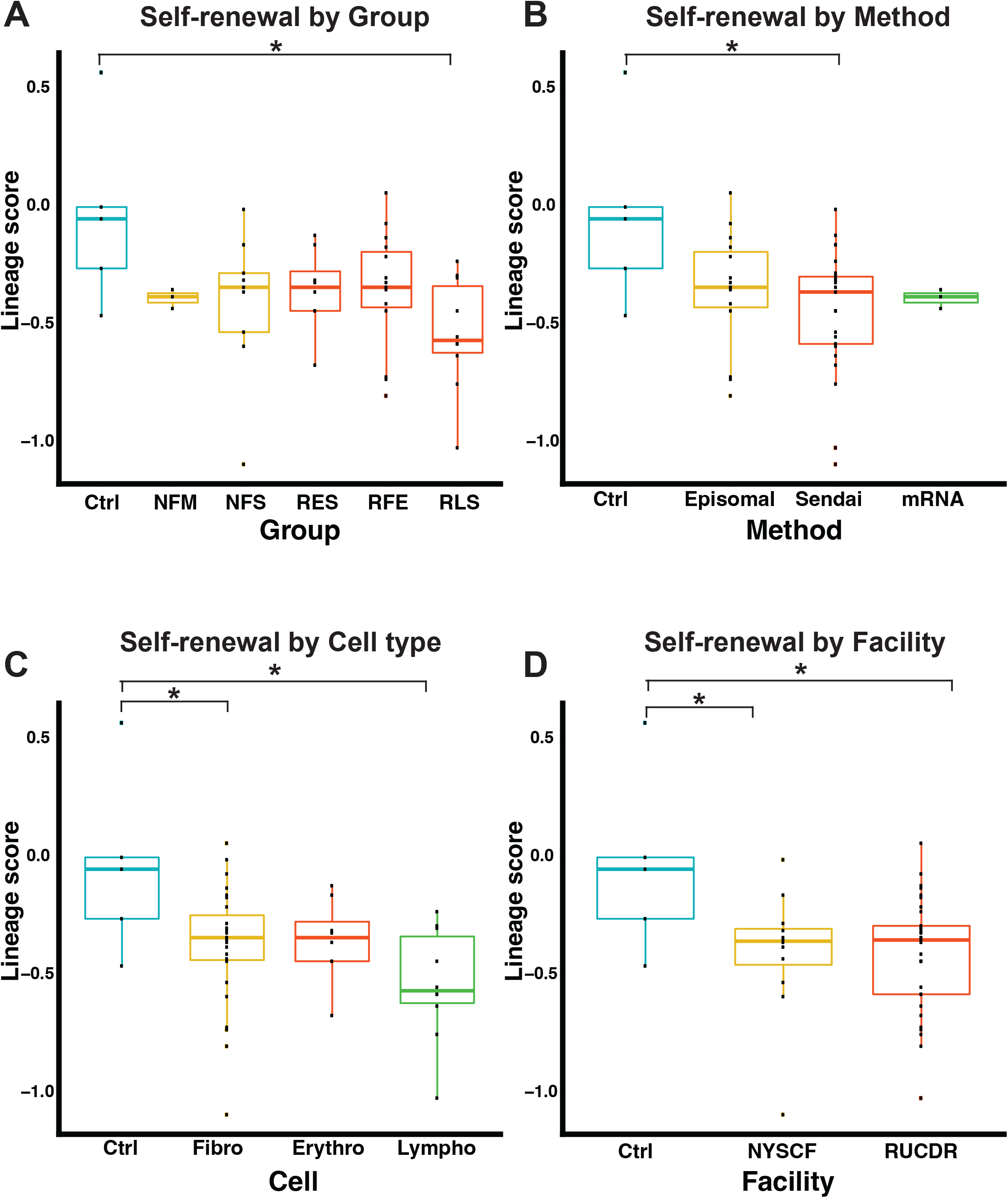
iPSC qPCR scorecard analysis. **A-D**, Self-renewal lineage scores from iPSCs analyzed by (A) reprogramming group, (B) factor delivery method, (C) starting cell type, and (D) reprogramming facility. One-way ANOVA. *p<0.05.

### 16p11.2del iPSCs display functional traits of pluripotency

While protein and transcript expression of pluripotency markers serves as an effective proxy for stem cell tri-lineage differentiation ability, we decided that more functional assessments were required to truly measure potential differences in reprogramming methods. We again utilized the TaqMan hPSC Scorecard panel towards this goal. Rather than using undifferentiated iPSC samples as input, we instead cultured each line as a three-dimensional embryoid body (EB) for 12 days to allow for spontaneous differentiation into the different germ layers. Given that our primary interest lies in the ability of these cells to become neurons, we focused our initial assessment on the ectodermal lineage. No significant differences were observed across the five 16p11.2del groups and the NT controls (Figure 4a). When the 16p11.2del iPSCs were grouped by their various characteristics, we only found a small but significant difference between patients 14758.x3 and 14799.x1 (Figure 4b-c, Supplemental Figure 3a-d).

**Figure 4:**
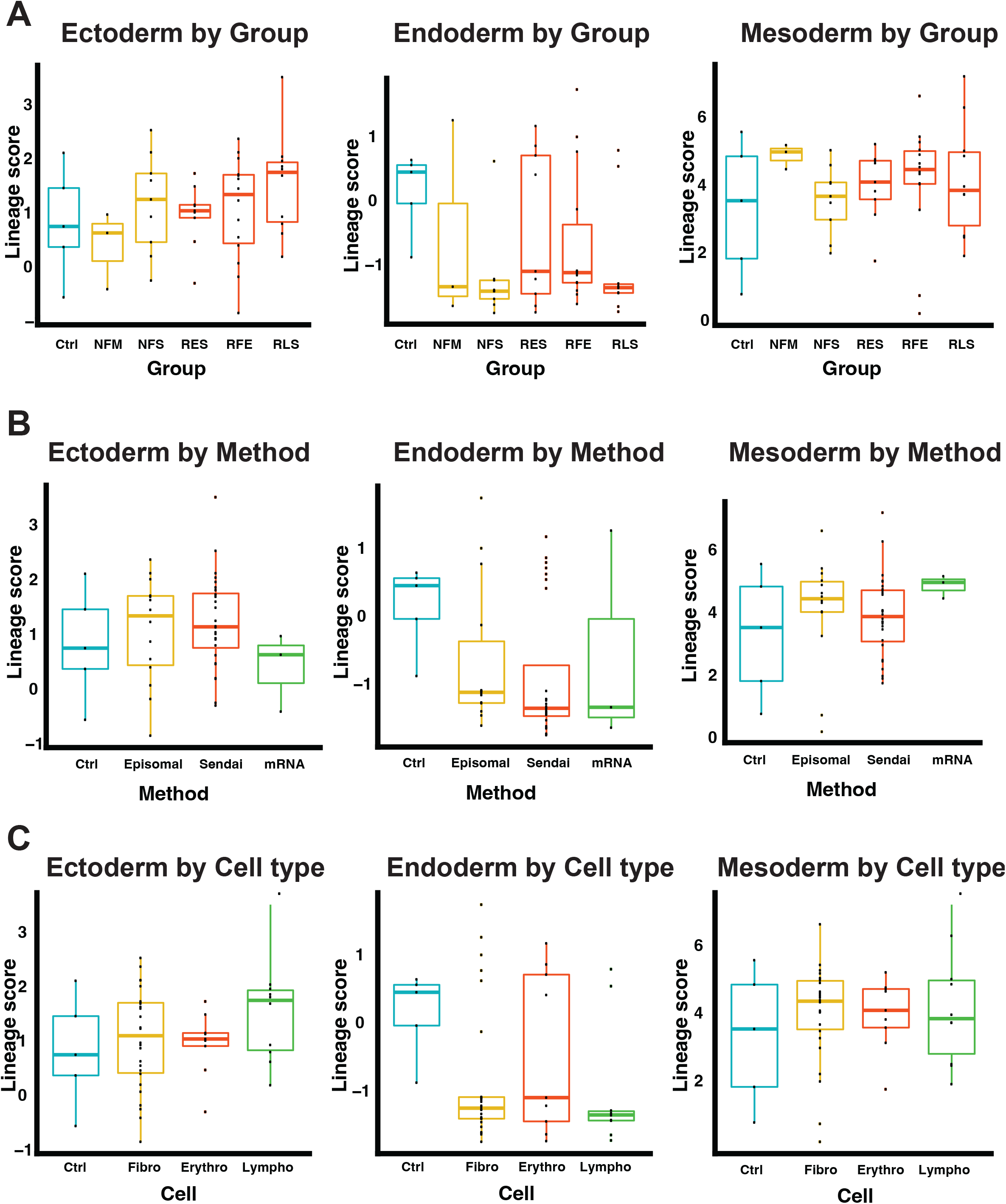
Embryoid body qPCR scorecard analysis. **A-C**, Ectoderm, endoderm, and mesoderm lineage scores from iPSC-derived embryoid bodies analyzed by (A) reprogramming group, (B) factor delivery method, and (C) starting cell type. One-way ANOVA. *p<0.05.

We then turned our attention to the results of the non-ectodermal lineage qPCR panels. We again found no differences among the five 16p11.2del groups in endoderm or mesoderm scores (Figure 4a). Our sub-categorization analysis did reveal a genotype-driven difference in endoderm score (Supplemental Figure 3a), though no other differences were observed (Figure 4b-c, Supplemental Figure 3b-d). Therefore, there does not appear to be a clear relationship between the five reprogramming groups and the observed lineage potential.

### 16p11.2del iPSCs differentiate into neurons

The 16p11.2del iPSCs appeared to be consistent in their ability to follow ectodermal trajectories, and as such, we were able to perform subsequent neuronal differentiations and comparative analysis on all 16p11.2del cell lines to determine the effects of reprogramming method on electrophysiological properties. We co-cultured both 16p11.2del and NT iPSC-derived neurons generated using doxycycline-inducible neurogenin2 (NGN2) overexpression with primary mouse glia in 12-well multi-electrode arrays (MEA). Over the next 6 weeks, we measured such neuronal firing properties as number of active electrodes, mean firing rate, and network burst rate. Grouped analysis of active electrodes and mean firing rate found no significant differences at any time across the five 16p11.2del cohorts, though the RFE neurons did show significant reductions in these parameters compared to NT controls by week 6 (Figure 5a). The RLS and RES neurons were also decreased in mean firing rate compared to NT controls at this time point. Importantly, while the 16p11.2del groups were similar to each other at week 6, all but the NFM neurons (p = 0.0567) displayed significantly impaired network burst frequency compared to the NT controls, suggesting that deletion of the 16p11.2 segment results in abnormal *in vitro* circuit development. It is possible the NFM did not reach significance due to the relatively smaller number of clones tested.

**Figure 5:**
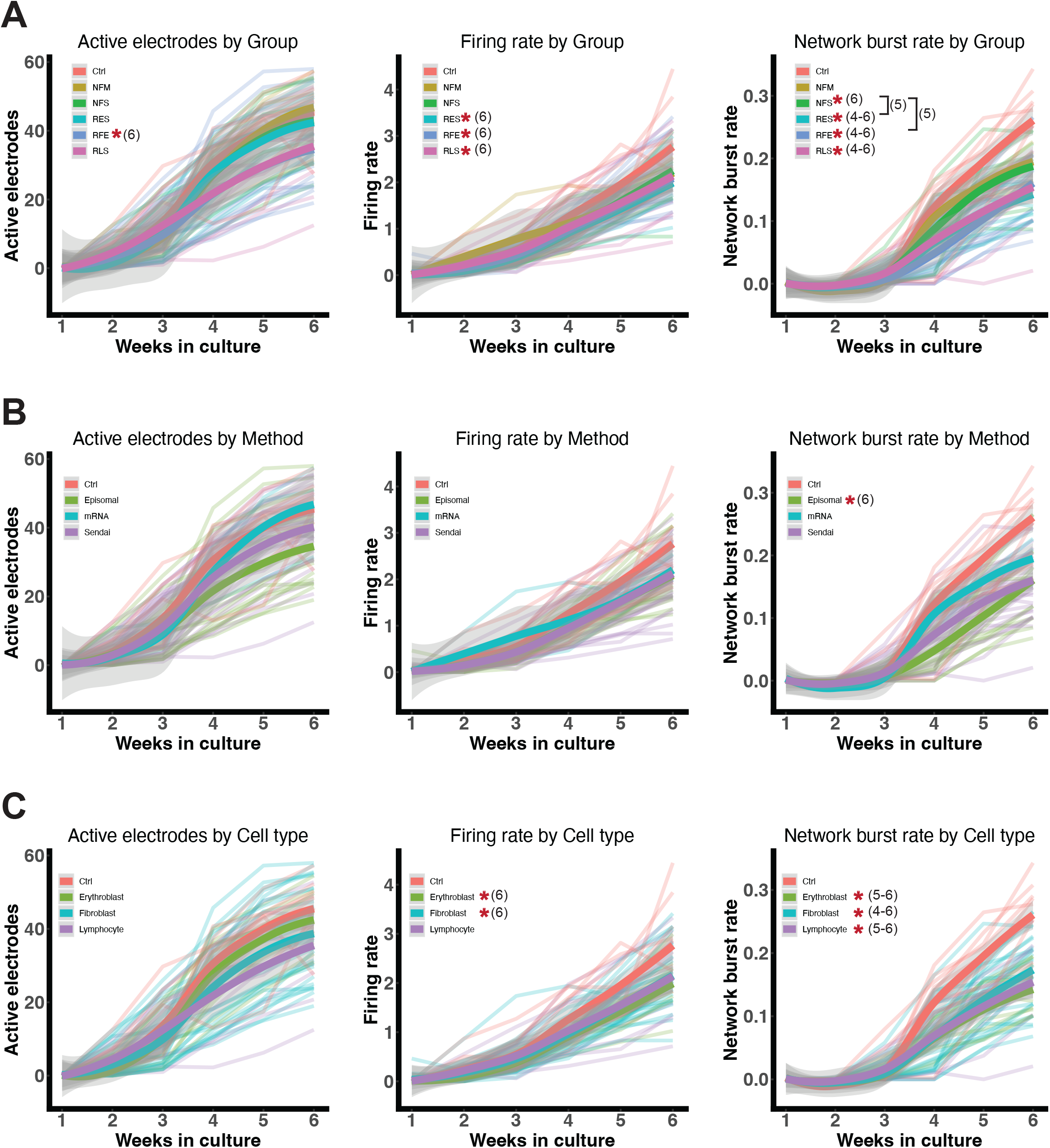
MEA analysis of iPSC-derived neurons. **A-C**, Number of active electrodes, firing rate, and network burst frequency of iPSC-derived neurons analyzed by (A) reprogramming group, (B) factor delivery method, and (C) starting cell type. Two-way RM ANOVA. *p<0.05 compared to controls. Number in parentheses refers to time point at which statistical significance reached.

We delved deeper into the MEA analysis by again sub-dividing the comparisons based on reprogramming method characteristics. Separating the cell lines by genotype again revealed significant differences in firing rate as early as week 5, and network burst frequency as early as week 4 (Supplemental Figure 4a). Provider comparison revealed no differences between RUCDR and NYSCF samples at week 6, though RUCDR neurons were reduced in firing rate and bursts compared to NT controls while NYSCF samples only showed reduced burst metrics at this time point (Supplemental Figure 4b). We observed no factor delivery method or starting cell type-mediated differences across the 16p11.2del neurons groups, though some comparisons with NT controls did reach significance (Figure 5b-c). Additionally, no statistically significant differences were observed in any of the reprogramming-based sub-comparisons of active electrode number, suggesting the reprogramming method does not impact the ability of iPSC-derived neurons to survive rigors of plating and 6 weeks in culture. Patient differences were found in firing rate and network bursting, perhaps hinting at variability in these parameters across the genetic background of 16p11.2del patients (Supplemental Figure 4c). Finally, grouping the data by donor sex revealed differences between male and female patient lines in firing rate and network burst rate (Supplemental Figure 4d). An important caveat to donor sex comparisons is the fact that there is only one female donor, so these results could be driven by the natural variation across patient lines described previously. Overall, we conclude that there are consistent genotype-driven differences in our MEA dataset, but that the overall impact of reprogramming methodology appears to be limited.

## DISCUSSION

There are many ways to assess iPSC reprogramming performance depending on the desired experimental endpoints. Given our interest in investigating human brain cells derived from patients afflicted by ASD, we based our assessment criteria on ectodermal and neuronal properties. We found no evidence that reprogramming methods consistently alter the pluripotency and neuronal differentiation abilities of their given iPSC products. The finding that reprogramming method appears to have no consequential effect on transcription factor-mediated directed neuronal differentiation has far-reaching implications for comparative *in vitro* studies of the brain that involve multiple iPSC lines generated using different methods from disparate sources, as our results suggest that inter-line differences are more likely to be driven by such factors as the genomic background of the donor^12,13^. As a result, high-throughput experiments utilizing dozens (if not hundreds) of stem cell lines from the many individual laboratories and institutional biobanks that are actively manufacturing iPSCs could be designed without great concern for differences in reprogramming practices.

This work is not without its shortcomings. First of all, the number of clones per cell line differed greatly which negatively impacted our ability to perform properly powered comparative analysis for some of the methods. Secondly, our neuronal differentiation experiments were based on NGN2-directed induction techniques, which have not been universally adopted. It is possible that iPSC-derived neurons generated using protocols that rely solely on small molecule inhibitors of key signaling pathways would not show the same consistency across reprogramming methods that we observed. Finally, when making our formal conclusions, we did not consider the potential for each reprogramming method to introduce genetic mutations not found in the donor somatic cells.

## ACKNOWLEDGEMENTS

This work was supported by a grant from the Simons Foundation (SFARI award 497451 to K.E. and A.M.)

---

**Supplemental Figure 1:**
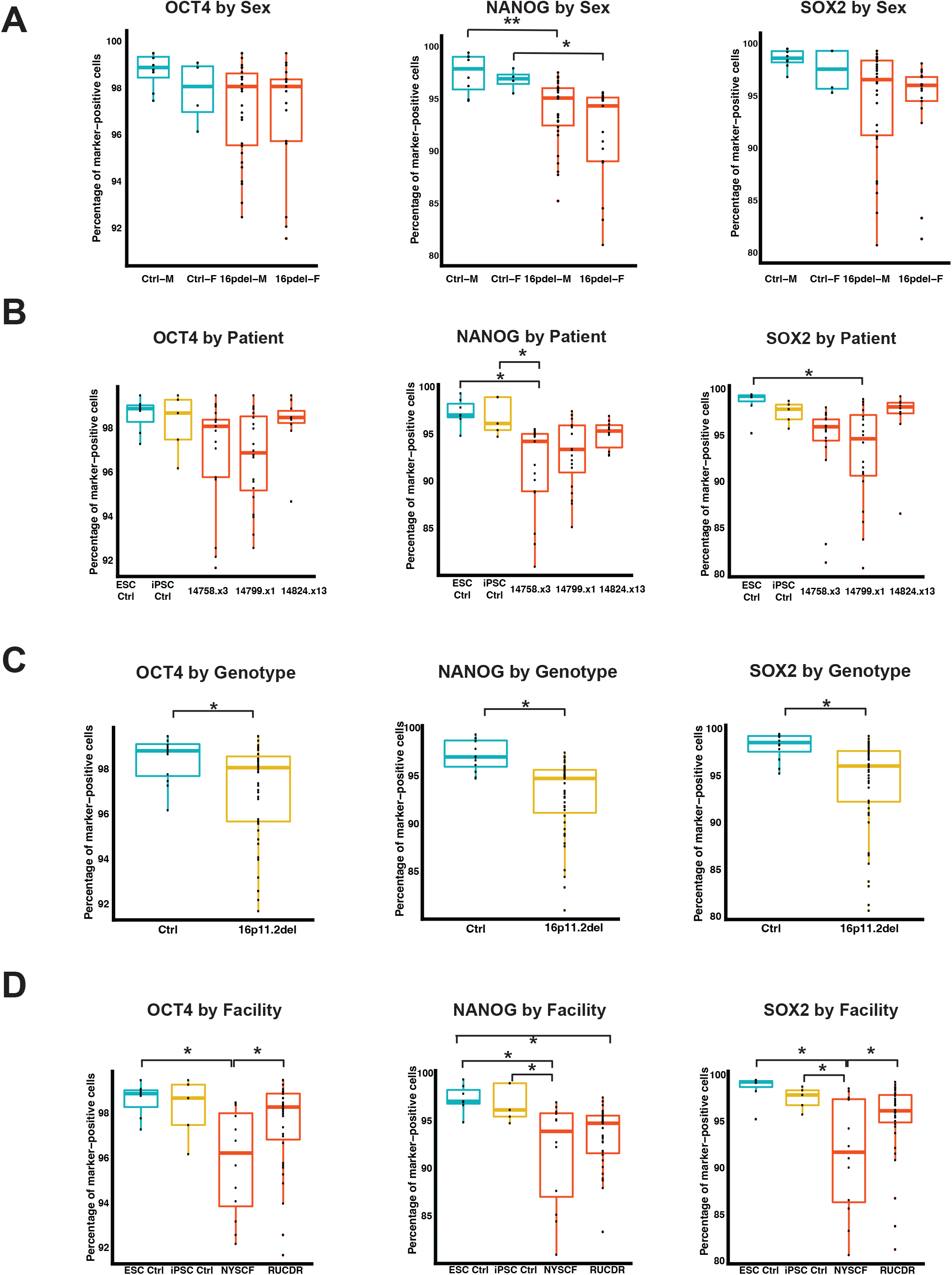
Additional iPSC FACS analysis. **A-C**, Percentage of OCT4-, NANOG-, and SOX2-positive cells as a function of (A) sex, (B) 16p11.2del patient, (C) genotype, and (D) reprogramming facility. One-way ANOVA. *p<0.05.

**Supplemental Figure 2:**
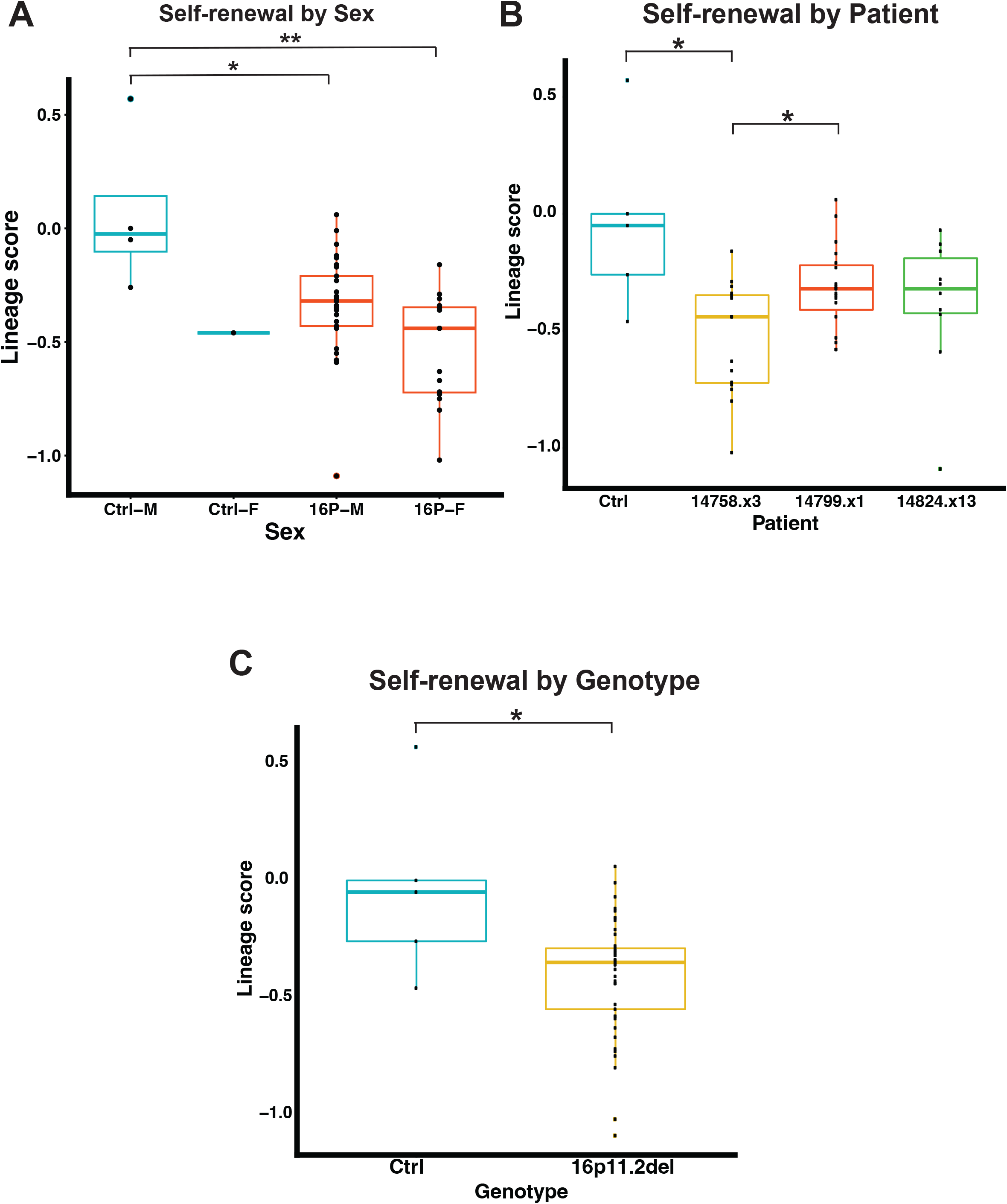
Additional iPSC qPCR scorecard analysis. **A-B**, Self-renewal lineage scores from iPSCs analyzed by (A) sex, (B) 16p11.2del patient, and (C) genotype. One-way ANOVA. *p<0.05.

**Supplemental Figure 3:**
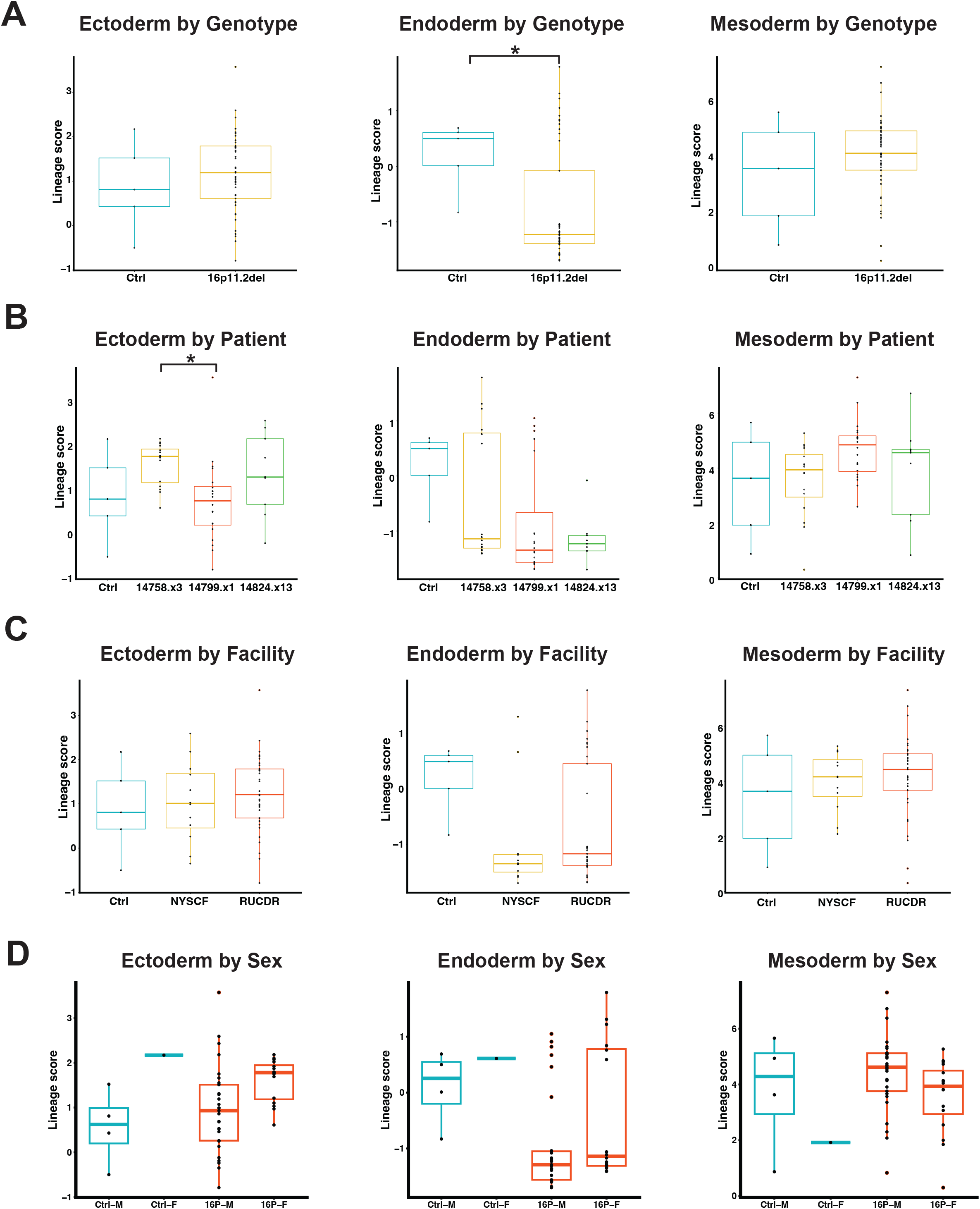
Additional embryoid body qPCR scorecard analysis. **A-C**, Ectoderm, endoderm, and mesoderm lineage scores from iPSC-derived embryoid bodies analyzed by (A) genotype, (B) 16p11.2del patient, (C) reprogramming facility, and (D) donor sex. One-way ANOVA. *p<0.05.

**Supplemental Figure 4:**
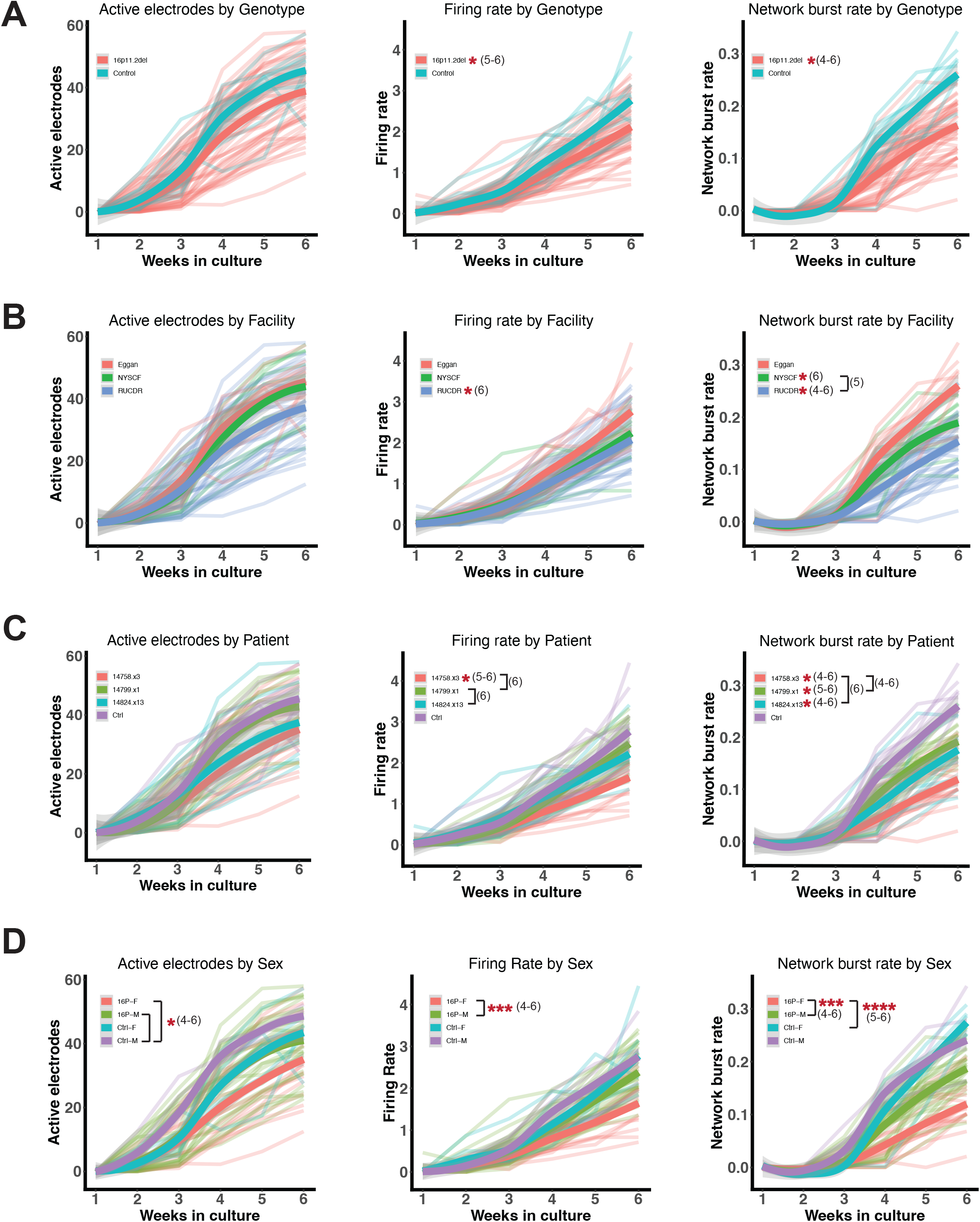
Additional MEA analysis of iPSC-derived neurons. **A-C**, Number of active electrodes, firing rate, and network burst frequency of iPSC-derived neurons analyzed by (A) genotype, (B) reprogramming facility, (C) 16p11.2del patient, and (D) donor sex. Two-way RM ANOVA. *p<0.05, ***p<0.001, ****p<0.0001. If no brackets shows, comparison is to controls. Number in parentheses refers to time point (week) at which statistical significance reached.

## The NYSCF Global Stem Cell Array Team

Lauren Bauer

Stephen Chang

Jordan Goldberg

Aana Kim Hahn

Premlatha Jagadeesan

Hesed Kim

Katie Krumholz

Gregory Lallos

Dorota Moroziewicz

Chris Napolitano Ana Sevilla

Keren A Weiss Chris

M Woodward Matthew Zimmer

## REFERENCES

1. Gao, S. et al. Unique features of mutations revealed by sequentially reprogrammed induced pluripotent stem cells. Nat. Commun. 6, 6318 (2015).

2. Lister, R. et al. Hotspots of aberrant epigenomic reprogramming in human induced pluripotent stem cells. Nature 471, 68–73 (2011).

3. Chang, G. et al. High-throughput sequencing reveals the disruption of methylation of imprinted gene in induced pluripotent stem cells. Cell Res. 24, 293–306 (2014).

4. Saha, K. & Jaenisch, R. Technical challenges in using human induced pluripotent stem cells to model disease. Cell Stem Cell 5, 584–595 (2009).

5. Vitale, A. M. et al. Variability in the generation of induced pluripotent stem cells: importance for disease modeling. Stem Cells Transl. Med. 1, 641–650 (2012).

6. Bock, C. et al. Reference maps of human es and ips cell variation enable high-throughput characterization of pluripotent cell lines. Cell 144, 439–452 (2011).

7. Silva, M. et al. Generating iPSCs: translating cell reprogramming science into scalable and robust biomanufacturing strategies. Cell Stem Cell 16, 13–17 (2015).

8. Weiss, L. A. et al. Association between Microdeletion and Microduplication at 16p11.2 and Autism. N. Engl. J. Med. 358, 667–675 (2008).

9. Qureshi, A. Y. et al. Opposing Brain Differences in 16p11.2 Deletion and Duplication Carriers. Journal of Neuroscience 34, 11199–11211 (2014).

10. McCarthy, S. E. et al. Microduplications of 16p11.2 are associated with schizophrenia. Nat. Genet. 41, 1223–1227 (2009).

11. Horev, G. et al. Dosage-dependent phenotypes in models of 16p11.2 lesions found in autism. Proceedings of the National Academy of Sciences 108, 17076–17081 (2011).

12. Wells, M. F. et al. Natural variation in gene expression and viral susceptibility revealed by neural progenitor cell villages. Cell Stem Cell 1–21 (2023).

13. Neavin, D. R. et al. A village in a dish model system for population-scale hiPSC studies. Nat. Commun. 14, 3240 (2023).

