## Supplemental Methods for "Reprogramming method does not impact the neuronal differentiation potential of 16p11.2 deletion patient iPSCs"

*Cell Line Donors*: All three cell line donors are participants in the Simons Searchlight (formerly known as Simons Variations in Individuals Project, Simons VIP[1]) study and donated fibroblasts and/or blood as part of the study protocol (Geisinger IRB #2011-0320). All donors are carriers of a copy number variant (CNV) deletion at 16p11.2 (16p11.2 del).

*Induced Pluripotent Stem Cell lines (iPSCs):* iPSCs were derived from donor fibroblasts, erythroblasts and/or lymphocytes by Rutgers University's Cell and DNA Repository (RUCDR, now Sampled) and the NYSCF, respectively. NYSCF reprogrammed fibroblasts using mRNA or Sendai Virus delivery of reprogramming factors as previously described[2,3]. RUCDR methods were as follows:

> Reprogramming of Erythroblasts. The reprogramming of erythroblasts relied on the selective culturing of the erythroblast population from either fresh PBMCs or from CPLs with the use of Erythropoietin, IGF-1, SCF, IL3 and ascorbic acid[4]. After expansion the erythroblasts were efficiently infected with Sendai viral vectors expressing Oct4, Sox2, Klf4 and C-Myc (CytoTune®-iPS 2.0 Sendai Reprogramming Kit from Life Technologies). Initial plating of erythroblasts undergoing reprogramming was done on irradiated mouse embryonic feeder cells. Multiplexed QPCR assays were used to detect rearrangements in the T cell receptor B and G loci[5] as a means of demonstrating heterogeneity in the clonal composition of iPSC colonies picked from a single plate of reprogrammed CPLs (Moore et al., unpublished).
>
> Reprogramming of Fibroblasts. iPSCs were obtained from fibroblasts using the Epi5™ Episomal iPSC Reprogramming Kit (Life Technologies) based on the transcription factors Oct4, Sox 2, Klf4, L-Myc, and Lin28[6], as well as a dominant negative p53[7].
>
> Reprogramming of CD4+ T cells. CD4+ T cells were isolated from either fresh or cryopreserved PBMCs using the CD4+ T Cell Isolation Kit (Miltenyi Biotech). Upon activation with CD3/CD28 beads (Dynabeads™ Human T-Activator CD3/CD28 for T Cell Expansion and Activation kit from ThermoFisher), these cells were infected with Sendai viral vectors expressing Oct4, Sox2, Klf4 and C-Myc (CytoTune®-iPS 2.0 Sendai Reprogramming Kit from Life Technologies). Reprogramming was done cells cultured on irradiated mouse embryonic feeder cells.

*Stem cell culture*: Human ESCs and iPSCs were maintained in mTeSR media (Stem Cell Technologies, 85850) on Geltrex basement membrane matrix (1:100; Life Technologies, A1413301). Cells were split every 4-5 days (when they reached 80-90% confluency) using a 15 minute/37°C incubation in Accutase (Innovative Cell Technologies, AT104) followed by 1:10 dilution in mTeSR. For each passage, media was supplemented with ROCK inhibitor Y-27632 (10 μM; Stemgent, 04-0012) for 24 hours after plating.

*Viral transduction & NGN2 neuron production:* TetO-Ngn2-Puromycin and Ubq-rtTA constructs were obtained from the Wernig lab (Stanford) before being packaged as high-titer lentiviruses (Alstem, Richmond, CA). When hPSCs reached 80-100% confluency, they were dissociated with Accutase before being re-suspended in lentivirus-containing mTeSR media supplemented with Y-27632 at a range of MOI = 1 to MOI = 3. Cells were then plated on Geltrex-coated 12-well plates at 500,000 cells per well in a total volume of 750 μl/well. After 18-24 hours, lentiviral media was aspirated and cells were fed with mTeSR media and maintained as described above. Transduced cells were maintained for up to 10 passages for inductions and transduction efficiencies typically ranged from 65-85% across cell lines. NGN2 neuron induction and maintenance was performed as previously described[8].

*Multi-electrode array (MEA)*: NGN2 neurons co-cultured with mouse glia were plated on a Geltrex-coated 12-well MEA plate (Axion Biosystems, M768-GL1-30Pt200) in Neurobasal complete media [Neurobasal media (97% v/v; Life Technologies 21103049), Glutamax (1:100), 20% Glucose (1.5% v/v), MEM-NEAA (1:200), B27 (1:50), BDNF (10 ng/mL), CTNF (10 ng/mL), and GDNF (10 ng/mL)] at 40,000 neurons/$cm^2$ + 70,000 glia/$cm^2$. Cells were fed 2-3 times per week with partial exchanges to reach a final volume consisting of 80% fresh media and 20% conditioned media. Five minutes of neuronal activity was measured weekly using the Maestro 12-well 64 electrodes per well micro-electrode array (MEA) plate system (Axion Biosystems, Atlanta, GA). Data was analyzed using the Axion Integrated Studio 2.4.2 and the Neural Metric Tool (Axion Biosystems).

*qPCR scorecard:* The embryoid body, qPCR-based scorecard of germ lineage differentiation potential was performed as previously described[9].

References


1. Spiro, J. E. & Chung, W. K. Simons Variation in Individuals Project (Simons VIP): A Genetics-First Approach to Studying Autism Spectrum and Related Neurodevelopmental Disorders. *Neuron* **73**, 1063–1067 (2012).
2. Paull, D. *et al.* Automated, high-throughput derivation, characterization and differentiation of induced pluripotent stem cells. *Nat. Methods* **12**, 885–892 (2015).
3. Zhou, H. *et al.* Rapid and efficient generation of transgene-free iPSC from a small volume of cryopreserved blood. *Stem Cell Rev.* **11**, 652–665 (2015).
4. Soares, F. A. C., Pedersen, R. A. & Vallier, L. Generation of human induced pluripotent stem cells from peripheral blood mononuclear cells using Sendai virus. *Methods Mol. Biol.* **1357**, 23–31 (2016).
5. Brüggemann, M. *et al.* Powerful strategy for polymerase chain reaction-based clonality assessment in T-cell malignancies Report of the BIOMED-2 Concerted Action BHM4 CT98-3936. *Leukemia* **21**, 215–221 (2007).
6. Yu, J., Chau, K. F., Vodyanik, M. A., Jiang, J. & Jiang, Y. Efficient feeder-free episomal reprogramming with small molecules. *PLoS One* **6**, e17557 (2011).
7. Hong, H. *et al.* Suppression of induced pluripotent stem cell generation by the p53-p21 pathway. *Nature* **460**, 1132–1135 (2009).
8. Nehme, R. *et al.* Combining NGN2 Programming with Developmental Patterning Generates Human Excitatory Neurons with NMDAR-Mediated Synaptic Transmission Resource Combining NGN2 Programming with Developmental Patterning Generates Human Excitatory Neurons with NMDAR-Mediated. *Cell Rep.* **23**, 2509–2523 (2018).
9. Tsankov, A. M. *et al.* A qPCR ScoreCard quantifies the differentiation potential of human pluripotent stem cells. *Nat. Biotechnol.* **33**, 1182–1192 (2015).
